# Clonal gametogenesis is triggered by intrinsic stimuli in the hybrid’s germ cells but is dependent on sex differentiation

**DOI:** 10.1101/2021.10.27.466081

**Authors:** Tomáš Tichopád, Roman Franěk, Marie Doležálková-Kaštánková, Dmitrij Dedukh, Anatolie Marta, Karel Halačka, Christoph Steinbach, Karel Janko, Martin Pšenička

## Abstract

Interspecific hybridization may trigger the transition from sexual reproduction to asexuality, but mechanistic reasons for such a change in a hybrid’s reproduction are poorly understood. Gametogenesis of many asexual hybrids involves a stage of premeiotic endoreduplication (PMER), when gonial cells duplicate chromosomes and subsequent meiotic divisions involve bivalents between identical copies, leading to production of clonal gametes. Here, we investigated the triggers of PMER and whether its induction is linked to intrinsic stimuli within a hybrid’s gonial cells or whether it is regulated by the surrounding gonadal tissue.

We investigated gametogenesis in the *Cobitis taenia* hybrid complex, which involves sexually reproducing species (*Cobitis elongatoides* and *C. taenia*) as well as their hybrids, where females reproduce clonally *via* PMER while males are sterile. We transplanted spermatogonial stem cells (SSCs) from *C. elongatoides* and triploid hybrid males into embryos of sexual species and of asexual hybrid females, respectively, and observed their development in an allospecific gonadal environment. Sexual SSCs underwent regular meiosis and produced normally reduced gametes when transplanted into clonal females. On the other hand, the hybrid’s SSCs lead to sterility when transplanted into sexual males, but maintained their ability to undergo asexual development (PMER) and production of clonal eggs, when transplanted into sexual females.

This suggests that asexual gametogenesis is under complex control when somatic gonadal tissue indirectly affects the execution of asexual development by determining the sexual differentiation of stem cells and once such cells develop to female phenotypes, hybrid germ cells trigger the PMER from their intrinsic signals.

**Significance Statement:** Although sexual reproduction is a dominant trait among all eukaryotes, many taxa have evolved the ability to reproduce asexually. While asexuality appears to be linked to interspecific hybridization, it remains unknown how the coexistence of diverged genomes may initiate such a swap in reproduction. In our study, we transplanted germ cells between asexual hybrids and their parents. On one hand, the ability of clonal gametogenesis occurred exclusively in hybrid germ cells, suggesting that asexual development is directly triggered by the hybrid genomic constitution of the cell. On the other hand, clonality was observed only in cells transplanted into females, suggesting that the execution of clonal development is influenced by signals from the gonadal environment and regulated by somatic factors.

## Introduction

Although sexual reproduction is a predominant characteristic of all multicellular eukaryotes, based on conserved molecular machinery controlling meiotic divisions, it has been disrupted many times. This has resulted in a variety of the so-called asexual reproductive modes that occur in most animal and plant phyla. Asexual lineages not only allow key questions about ultimate advantages and disadvantages of sex to be tested, but due to their clonal multiplication, unusual gametogenic and developmental pathways, they have proved to be appealing models for many biological disciplines concerned with ecology, cell biology and molecular genetics (1). Yet, despite intensive research in asexual organisms, frustratingly large gaps remain in our understanding of mechanisms that trigger such a switch from sex to various forms of asexuality. This is apart from some straightforward cases, such as *Wolbachia*-induced asexuality (2) or the presence of candidate “asexuality genes” in a few model taxa (3, 4).

A promising class of theories aiming to identify some common mechanisms underlying the emergence of asexuality builds on the fact that many asexual organisms are of hybrid origin. It has been proposed that abandonment of sex may be stimulated by aberrant interactions between orthologous copies of individual genes (5, 6), chromosomes (7) or even entire regulatory networks brought together by hybridization between distinct but not co-adapted genomes (8). Unfortunately, the scarcity of empirical studies prevents any clear-cut conclusions about the role of hybridization in triggering asexuality. Indeed, if hybridization is supposed to initiate asexuality, it is difficult to explain why meiosis is affected in similar ways across diverse taxa.

For instance, one gametogenic pathway that is relatively common among independently arisen asexual animals and plants in *sensu lato* is premeiotic endoreplication (PMER) (9–16). This pathway, depicted in Fig. 1, is characterized by duplication of chromosomes in oogonia before meiosis. Consequently, subsequent meiotic divisions occur in a polyploid gamete with bivalents forming between homologues. Such a process alleviates potential problems in pairing of orthologous chromosomes in hybrids (9, 17) and simultaneously leads to the production of clonal progeny because bivalent pairing and crossovers occur between identical sister copies of duplicated chromosomes.

**Figure 1.**
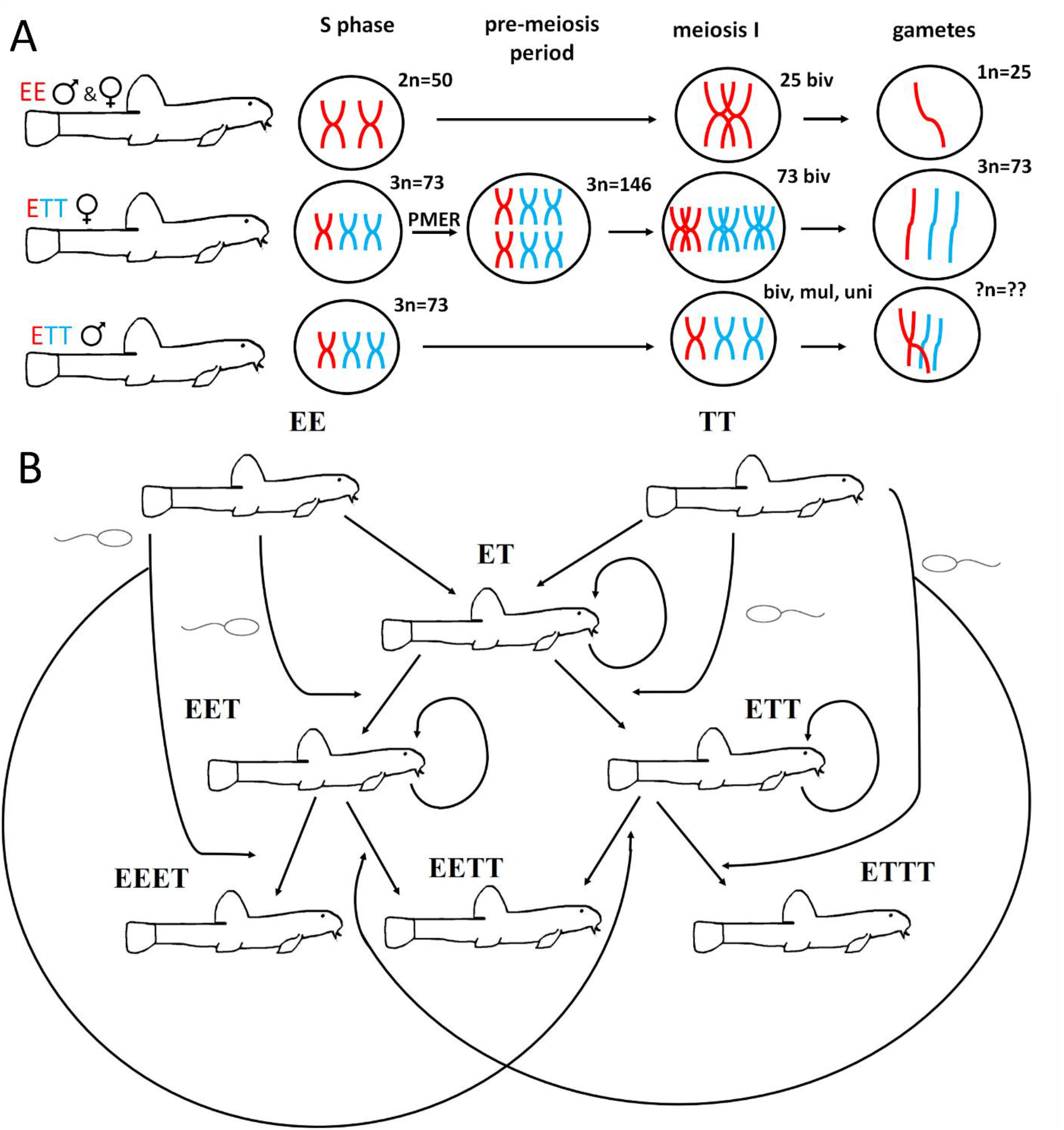
*Cobitis taenia* (TT) and *Cobitis elongatoides* (EE) hybridization scheme. A) Gametogenesis in EE, triploid ETT female and male. Red color represents chromosomes derived from parental species EE, blue those from parental species TT. In hybrid females, the premeiotic endoreplication (PMER) results in doubling of chromosomes and pairing between identical copy produces proper bivalents with crossovers. However, in males, PMER does not occur. Therefore chromosomes cannot properly pair leading to bivalents (biv), multivalents (mul) and univalents (uni). Parental species produce haploid gametes, hybrid females produce clonal eggs and males either cannot finish meiosis properly or final spermatozoa are often aneuploid or polyploid with disrupted motility. B) F1 hybrids are produced by natural or artificial spawning between TT and EE individuals. ET hybrid males are sterile while hybrid females are fertile using gynogenesis as reproductive mode. In some cases, sperm from either parental species can fertilize the egg giving rise to the triploid hybrids (ETT or EET depending on sperm), which are also either sterile (males) or fertile (females). Fertilization of triploid eggs is also possible but natural occurrence of EEET, EETT, and ETTT is very rare and tetraploids appear unable to reproduce.

Analysis of the speciation process in spined loaches (*Cobitis*; Actinopterygii) demonstrated that emergence of hybrid asexuality mechanistically coincided with hybrid sterility (9, 18). Hence, the emergence of asexual gametogenesis in general, and PMER in particular, may represent an alternative type of reproductive incompatibility promoting speciation among hybridizing species (18, 19). There are notable analogies between the emergence of hybrid asexuality and classical speciation models assuming the accumulation of postzygotic reproductive incompatibilities. Indeed, the emergence of both hybrid asexuality and sterility effectively restricts interspecific geneflow thereby promoting speciation, and scale with genomic divergence between parental species. Furthermore, in a similar way to hybrid sterility, asexual gametogenesis also arises asymmetrically with respect to the sex of the hybrids so that most known asexual vertebrates exhibit strongly female-biased sex ratios for which they are sometimes also referred to as ‘all-female’ or ‘unisexual’ (20, 21).

Unfortunately, there are only a few studies investigating functional differences between male and female hybrids in taxa where asexuality occurs. Available data indicate that when hybrid females are fertile and asexual (often employing PMER), their hybrid brothers are usually sterile (11, 19, 22). However, some exceptions exist, *e*.*g*. in hybridogenetic taxa like *Pelophylax* (23) or *Squalius* (24). Crossing experiments within several asexual complexes of fishes and reptiles further demonstrate that asymmetry between female hybrid asexuality and male sterility is directly linked to the merging of parental genomes and already occurs in the F1 hybrid generation (9, 11, 25–27). Yoshikawa *et al*. (28) further examined *Misgurnus* spp. (loaches), where hybrid females typically reproduced clonally with PMER, and artificially reverted clonal diploid female progenies into males. They reported that PMER occurred also in spermatogonia of these artificially sex-reverted males from females, which was surprising since natural *Misgurnus* male hybrids are sterile (11). Such a finding therefore indicates that asexual gametogenesis is somehow linked to one sex (females), and seems to depend on genetic sex determination rather than on phenotypic sex.

PMER thus appears to be a crucial cellular deviation in the evolution of many natural hybrid and allopolyploid lineages, making it important in furthering our understanding of genetic and cellular mechanisms which underly its occurrence. Unfortunately, how and why the hybrid germ cells switch their developmental pathway towards PMER are unknown. For instance, it remains unclear whether PMER results from endomitosis, which involves mitotic replication of chromosomes without cell division, or endoreduplication, *i*.*e*. replication of chromosomes without initiation of mitosis. It has been shown that red crucian carp × common carp [*Carassius auratus* (red variety) x *Cyprinus carpio*] hybrids produce tetraploid oogonia by germ cell fusion, rather than by multiplication of their chromosomes (29).

Loaches of the family Cobitidae (Cypriniformes, Teleostei) have been shown to be a suitable model organism to understand the mechanisms underlying hybrid sterility and asexuality (10, 17, 18, 28, 30). In particular, the so-called *C taenia* hybrid complex is distributed in Europe and comprises sexual species of *Cobitis taenia* (TT) and *Cobitis elongatoides* (EE) that diverged ∼9 million years ago (18) but are frequently hybridizing, producing sterile males and hybrid females, which reproduce clonally using PMER (9, 31). Hybrid females are gynogenetic and hence sperm is required to trigger their gametes’ development but this does not generally contribute to the progeny’s genome. However, their oocytes sometimes incorporate the sperm’s genome leading to a new generation of sterile triploid males and gynogenetic females with a triploid genome composition (Figure 1). Consequently, natural populations of spined loaches are generally composed of sexual host species (often occurring in a minority), diploid, and mainly triploid clonal hybrid females (32).

In this study, we investigated whether the initiation of PMER is autonomously regulated in the hybrid’s germ cells or whether it depends on extrinsic stimuli from surrounding somatic cells and the tissue in which they occur. We also tested whether PMER is strictly confined to female sex determination, or whether the germline originating from males may also undergo such a pathway. To do so, we performed the following study, the design of which is depicted in Figure 2. We transplanted testicular cells containing spermatogonial stem cells (SSCs) between sexually reproducing species and their asexual hybrids from the *C. taenia* hybrid complex and investigated the development of such cells in the host’s body. Specifically, we extracted spermatogonial stem cells from *C. elongatoides* males (sexual species) and sterile allotriploid males with the *Cobitis elongatoides-taenia-taenia* genomic constitution (see Figure 1 for an explanation of the hybrid origins). These cells were reciprocally transplanted into juvenile recipients of both sexes that were sterilized by oligonucleotide morpholino treatment prior to transplantation. Therefore, two groups of fish were obtained: 1) triploid recipients of gonial cells from diploid donors (hereafter called TrDd) and 2) diploid recipients of gonial cells from triploid donors (hereafter called DrTd). Recipients were kept until adulthood and allowed to spawn in order to investigate their fertility and inheritance patterns in their progeny. After spawning, the recipients’ gonads were investigated using cytogenetic methods to check for meiotic patterns and potential presence of PMER.

**Figure 2.**
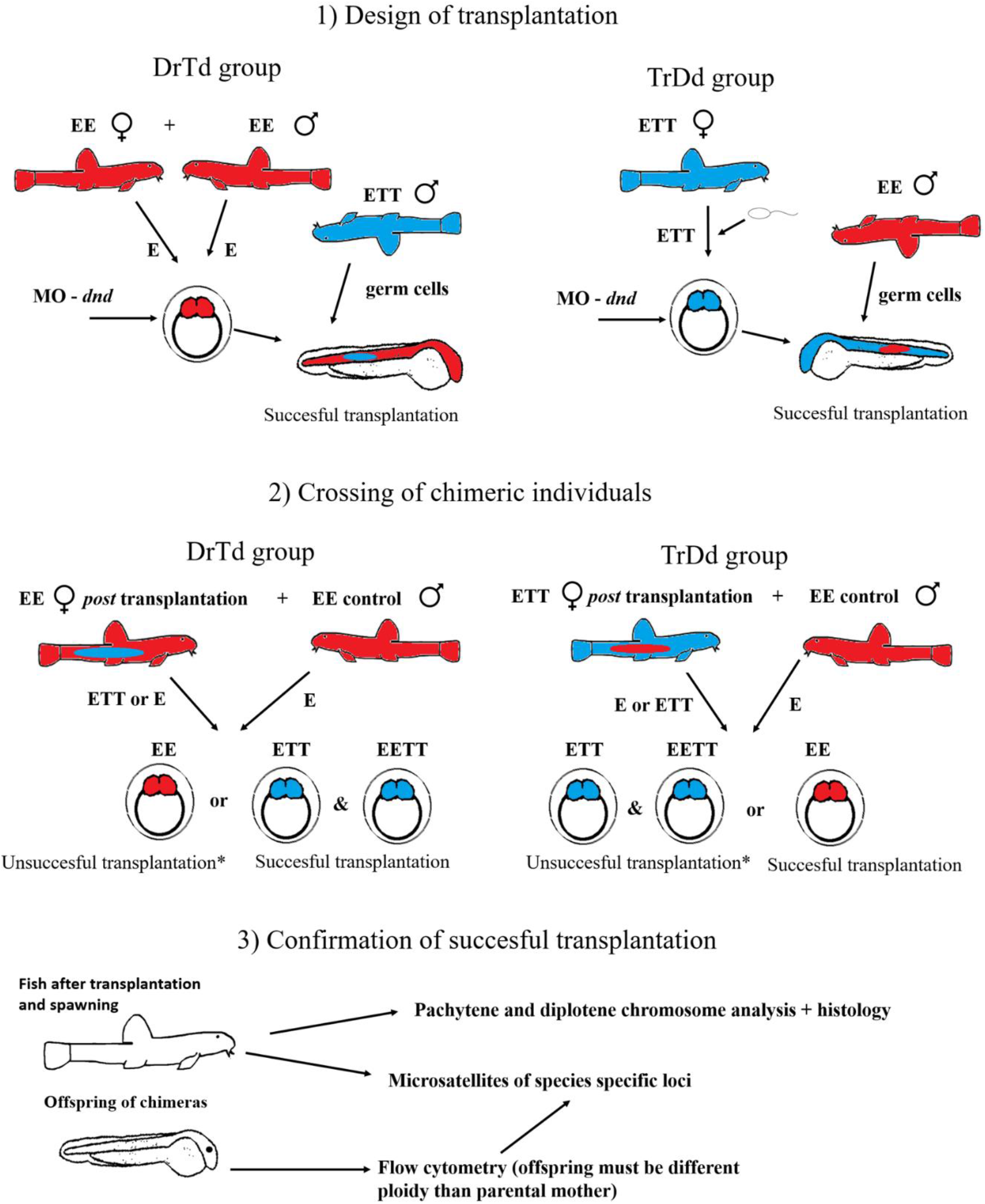
The experimental design. 1) The design of reciprocal transplantation between two groups: diploid recipient and triploid donor (DrTd), and triploid recipient and diploid donor (TrDd). In the DrTd group, parental species of *Cobitis elongatoides* (EE – red color) were spawned and their early embryos (2 cells stage) were injected with morpholino (MO) against the *dnd* gene to terminate development of parental gonads. Transplantation was undertaken using the germ stem cells from adult allotriploid male *Cobitis elongatoides-taenia-taenia* (ETT – blue color). In the second group TrDd, eggs of ETT females were activated with goldfish (*Carassius auratus*) (sperm symbol). Embryos were treated with anti-*dnd* MO and later transplanted with germ cells from adult EE males. 2) Two years after transplantation, experimental fish from both groups were spawned with the EE males. In the DrTd group, EE fish after successful transplantation should produce triploid ETT eggs where two scenarios can occur: either eggs are activated only (gynogenesis) producing ETT offspring or sperm can be incorporated into the eggs thus producing EETT. In the TrDd group, successfully transplanted ETT fish should produce haploid E gamete which must be fertilized with sperm and produce parental species EE. *In the case of unsuccessful morpholino treatment and transplantation, fish would produce their natural biotype, *i*.*e*. haploid eggs in the case of parental species and triploid eggs in case of hybrids. 3) Confirmation of successful transplantation. Offspring of chimeric fish from the DrTd group carried the *Cobitis taenia* microsatellite loci in the case of successful transplantation. On the other hand, *C. taenia* loci were absent in offspring from the TrDd group.

## Results

### Transplantation efficiency

Morpholino was injected into embryos between the one and four cell stage of development. After 2 years, we collected 31 representative fish of both experimental groups for further examinations, which included experimental crossing, histology and cytogenetic analysis of meiotic chromosomes. In total, we examined 23 fish of the DrTd group (12 females and 11 males) and eight fish of the TrDd group. Among these, we observed that four out of 12 DrTd females were not sterilized completely due to morpholino failure, six were sterilized but without successful transplantation (Figure 3) and two appeared as chimaeras after successful sterilization and transplantation. Similarly, sterilization failed in four out of 11 DrTd males. In five we observed sterilization without transplantations (Figure 3) and two appeared as successful chimaeras. In the TrDd group, six hybrid females were successfully sterilized and transplanted (Table 1) while two were succesfully sterilized but with no success in transplantation

**Figure 3.**
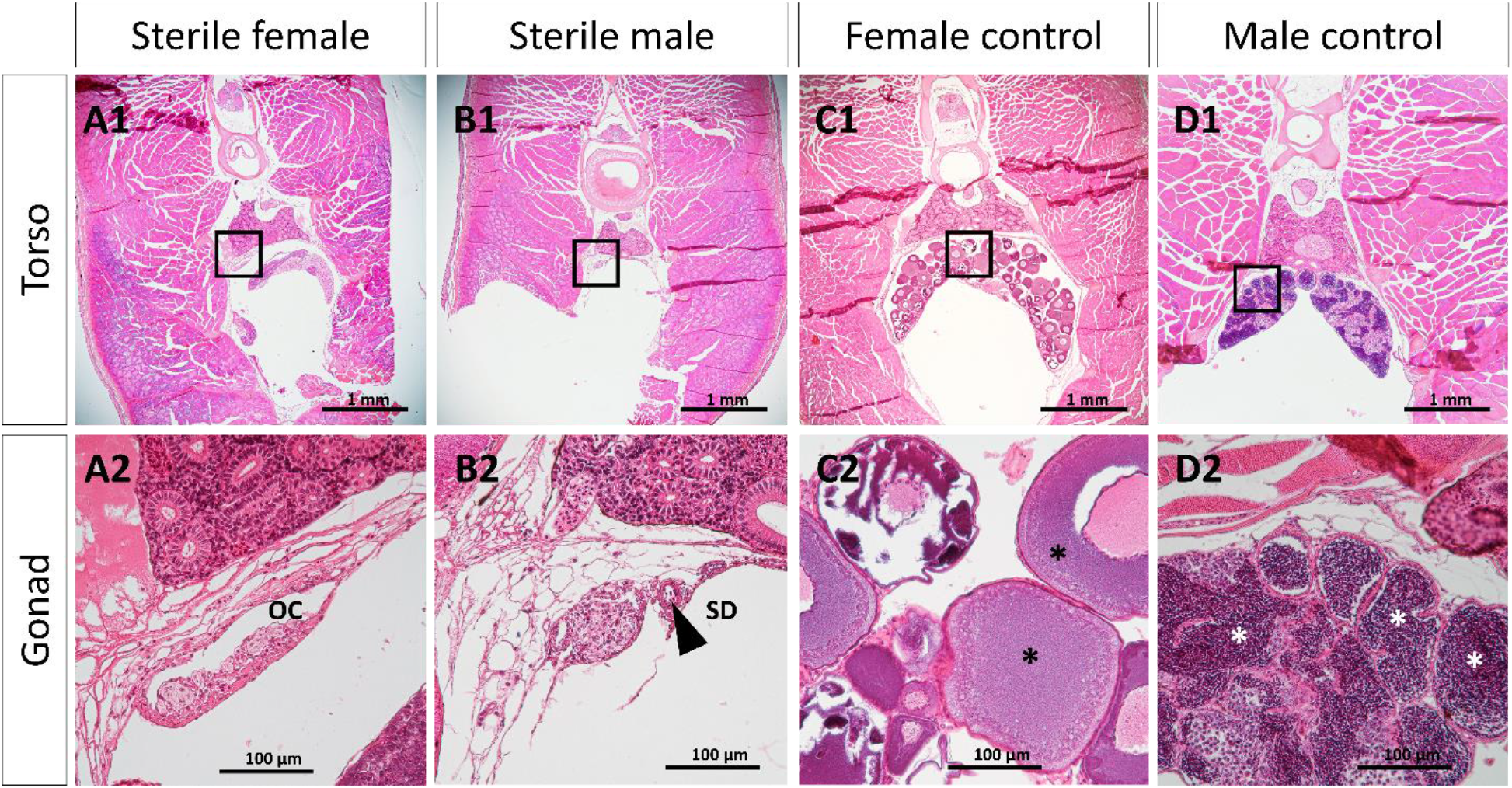
Results of histology analysis after MO-*dnd* treatment designed for *Cobitis taenia* in *Cobitis elongatoides*. A1 shows the whole body histology of a sterile female after morpholino treatment. A2 is a magnification of the ovarian cavity. B1 shows the whole body histology of a sterile male after morpholino treatment. B2 shows enlarged fragment from B1 of the sperm duct (SD), *i*.*e*. the connection between each testis to a urogenital opening, the testes themselves are not present. C1 and C2 show fertile females with vitellogenic eggs (black asterisks), while D1 and D1 show fertile males with gametes (white asterisks).

**Table 1.** Summary of the chimeric experiment. DrTd group, diploid recipient, triploid donor; TrDd group, triploid recipient, diploid donor; Biotype EE, parental species diploid *Cobitis elongatodies*; ETT, triploid hybrid of *Cobitis elongatodies-taenia-taenia*. Genetic profiling is based on microsatellite loci from chimeric offspring where the first number represents embryos derived from the recipient germ line and the second number from the donor germ line (*i*.*e*. successful transplantation). Note that the term “ordinary” in the histology analysis column stands for no difference from the control groups. We show only successful chimeric fish.

| ID | Sex of recipient | Biotype of recipient | Biotype of donor | Genetic profiling of offspring (donor derived/recipient derived) | Pachytene analysis | Histology analysis |
| --- | --- | --- | --- | --- | --- | --- |
| **DrTd group** |  |  |  |  |  |  |
| **5** | F | EE | ETT | ETT 16 & EE 0 | 73 bivalents | ordinary gonad with eggs |
| **6** | F | EE | ETT | ETT 7 & EE 0 | not done | NA |
| **13** | M | EE | ETT | not spawned | bivalents with univalents | reduced number of germ cells: no spermatozoa |
| **16** | M | EE | ETT | now spawned | bivalents and univalents | reduced number of germ cells:<br>no spermatozoa |
| **TrDd group** |  |  |  |  |  |  |
| **17** | F | ETT | EE | not spawned | 25 bivalents | ordinary gonad with eggs |
| **18** | F | ETT | EE | EE 8 & 0 ETT | 25 bivalents | ordinary gonad with eggs |
| 19 | F | ETT | EE | EE 7 & 0 ETT | 25 bivalents | ordinary gonad with eggs |
| 20 | F | ETT | EE | EE 17 & 0 ETT | not done | NA |
| 21 | F | ETT | EE | EE 15 & 0 ETT | not done | ordinary gonad with eggs |
| 22 | F | ETT | EE | Not spawned | 25 bivalents | NA |

### Genetic profiling of experimental fish and their offspring based on microsatellite markers

After reaching sexual maturity, 23 of the recipient fish were spawned with males of *C. elongatoides* and their offspring were profiled together with their parents. Genetic profiling of fish was performed by analyses of selected species specific (*C. elongatoides v. C. taenia*) microsatellite loci. Since we used two types of fish, diploid *C. elongatoides* and triploid *C. elongatoides-taenia-taenia*, the success of transplantation was indicated by the presence of *C. taenia* specific loci in the juveniles of the DrTd group analyzed (where parental species *C. elongatoides* produce juveniles with *C. taenia* loci) or by its absence in juveniles of TrDd (where hybrids *C. elongatoides-taenia-taenia* produce juveniles without *C. taenia* loci). Altogether, from 10 potential chimeric females (six *C. elongatoides* and four *C. elongatoides-taenia-taenia*), we obtained 136 juvenile individuals for analysis. Successful transplantation was confirmed in two out of six diploid *C. elongatoides* females of the DrTd group, which altogether produced 23 juveniles with a foreign allelic profile corresponding to the donor’s genotype (ETT). Moreover, in six out these 23 juveniles, we observed incorporation of paternal *C. elongatoides* sperm, leading to an increased ploidy level of progeny (EETT). This confirms their successful transplantation and demonstrates that such transplanted oogonia maintain their ability to produce clonal progenies. In the remaining four *C. elongatoides*, transplantation was not successful. In a reciprocal experiment, four *C. elongatoides-taenia-taenia* females from the TrDd group yielded 47 juveniles, whose genotype corresponded to one haploid set of their *C. elongatoides* donor and a second haploid set of the *C. elongatoides* father. This indicated successful transplantation in the TrDd group and suggested that *C. elongatoides* germ cells transplanted into ETT females conserved the ability to properly divide into apparently normal EE-type gametes.

### Histology

Histological analysis of chimeric fish was mostly directed to the males. Hybrid male sterility in *C. elongatodies x C. taenia* was represented by meiotic arrest leading to an aberrant germ cell population, lacking functional spermatozoa, while hybrid females produced fertile eggs. Nevertheless, it was possible to obtain supporting evidence of successful transplantation in diploid males (Figure 4A) when compared to gonadal tissue from diploid (Figure 4B) and triploid controls (Figure 4C). The DrTd males and triploid control males contained germ cells in premeiotic stages and a small number of postmeiotic abnormal cells, while diploid controls contained mainly functional spermatozoa. In females, diploid controls, DrTd and TrDd groups showed normal gametogenesis with late vitellogenic oocytes. However, histology itself was not used as an indicator of successful transplantation.

**Figure 4.**
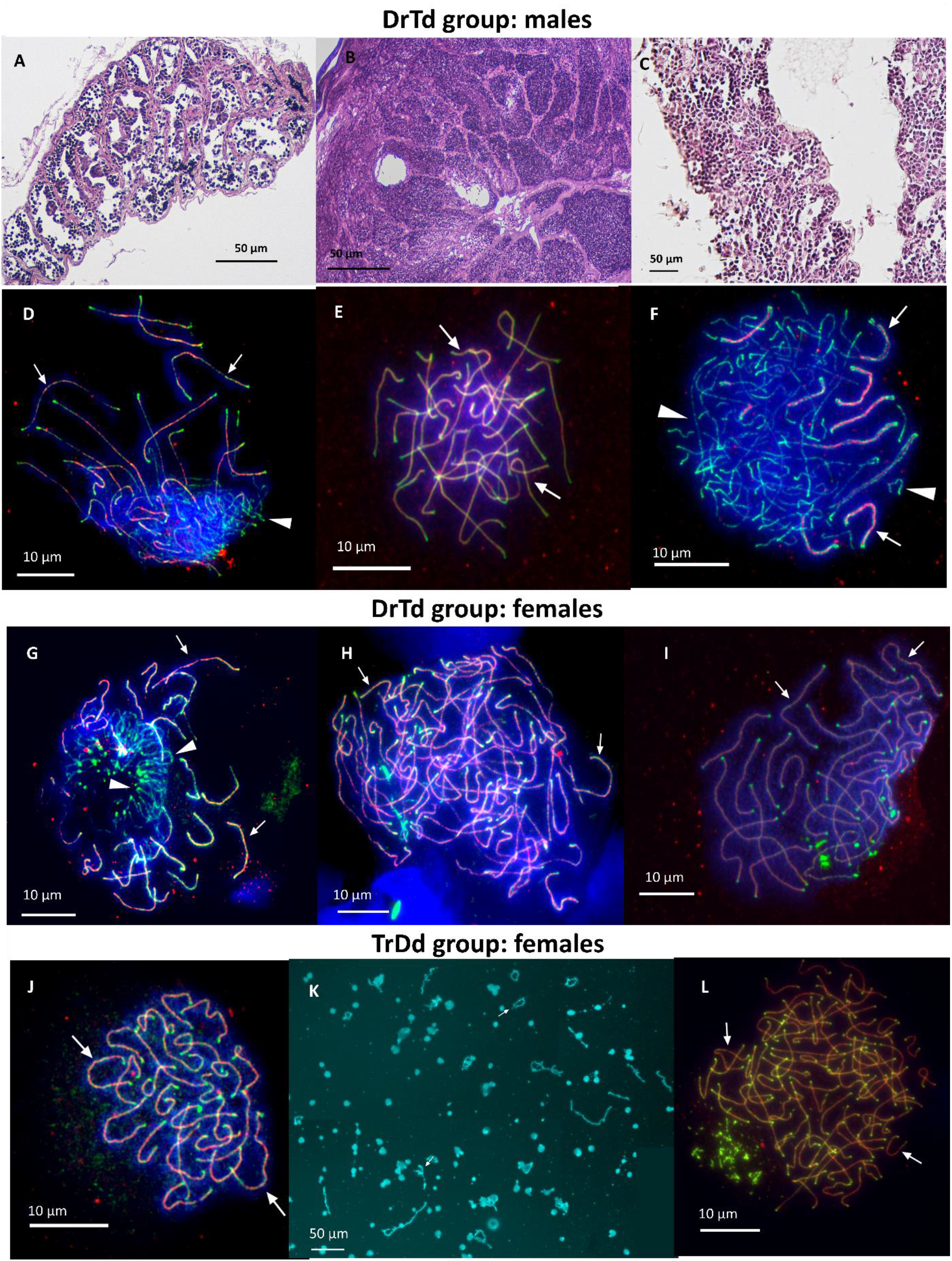
Results of histology and pachytene analysis from a diploid recipient with a triploid donor (DrTd, figures A-I), and a triploid recipient with a diploid donor (TrDd, figures J-L). A-C shows comparisons of histology of diploid chimeric fish (A) with a diploid control (B) and a triploid control (C). D-F show the pachytene analysis from the same fish samples. Diploid chimeric male (D) had an improper pairing during the pachytene stage leading to many univalents, while the diploid control (F) shows 25 bivalents in proper pairing. The triploid male control also had improper pairing which is similar to the diploid chimeric fish. Pachytene analysis of the female in the DrTd group shows a phenomenon typical of triploid females described by Dedukh (9, 40), *i*.*e*. either improper pairing (G) or properly paired 74 bivalents as result of premeiotic endoreplication (H). The diploid female control (I) shows 25 bivalents. (J) and (K) show the pachytene and diplotene chromosome analysis with 25 bivalents, respectively, in triploid fish from the TrDd group. Synaptonemal complexes were immunolabeled with antibodies against SYCP3 protein (green) and SYCP1 protein (red); chromosomes are stained with DAPI (blue). In K microphotograph, diplotene chromosomes are stained with DAPI (cyan).

### Meiosis analysis of chimeric fish

In order to investigate the particular gametogenic mechanisms adopted by the fish and their gametes, we isolated chromosomes during the pachytene and diplotene meiosis stages. We analyzed 67 pachytene spreads obtained from two *C. elongatoides* sperm lacking males in the DrTd group. We observed abnormal pairing with several bivalents and univalents (Figure 4D) and compared these with controls from 2n and F1 3n males (Figure 4E and 4F, respectively). We did not observe any cells with duplicated genomes. We additionally checked another six males from the same group; however, we did observe cells with 25 bivalents during pachytene, suggesting no success in transplantation. In females from the DrTd group, we did observe oocytes with improper pairing (Figure 4I) but also with duplicated genomes as 73 bivalents indicating the occurrence of PMER in chimeric fish during gametogenesis (Figure 4J). Figure 4K shows pachytene chromosomes from the 2n female controls. We did not find mispaired chromosomes in cells which underwent PMER. From four chimeric triploid females in the DdTr group, we successfully observed diplotene chromosomes with 25 bivalents (Figure 4M), and for one female we additionally managed to obtain chromosomes during both the pachytene and diplotene stages. In total, we examined 61 oocytes which included 25 bivalents of *C. elongatoides*. This suggests successful transplantation followed by the normal formation of haploid gametes of *C. elongatoides*. Triploid control fish can be seen in Figure 4N with 73 bivalents.

## Discussion

Although asexual organisms are important models for many biological disciplines, the mechanisms triggering clonal reproduction of gametes remain generally unclear. To test whether stimuli for asexual development are intrinsic to differentiating germ cells or depend on their gonadal environment, we performed reciprocal transplantation of spermatogonial cells between triploid hybrids (*C. elongatoides-taenia-taenia*) and one of their parental sexual species (*C. elongatoides*).

Prior to the experimental transplantation of the donor’s germ cells, the recipients’ gonads had to be sterilized. This goal was achieved with morpholino antisense RNA against the *Dnd* gene, which is a routinely applied procedure in many cell transplantation experiments. This is quite challenging when employed in hybrids and allopolyploids, because of the need to successfully target all gene copies coming from diverged parental subgenomes in a hybrid (33). Nevertheless, although *Cobitis* hybrids combine genomes that diverged as long as ∼9 Mya (18), we achieved a reasonable rate of successful sterilization ranging between 65-100 %, which is comparable to other studies (34, 35) and indicated that our MO design based on one parental genome is sufficient to targets all orthologous alleles in such situations.

After their successful transplantation, we recognized that male germ stem cells from male donors, either *C. elongatoides* or the triploid hybrid, transdifferentiated into oogonia when transplanted into female recipients. This demonstrated that both types of male germ stem cells were sensitive to the gonadal environment and their sexual differentiation is largely driven by the recipient’s body. This observation is consistent with a hypothesis that fish germ stem cells are sexually plastic (so-called sexually bipotent) and after transplantation, they may transdifferentiate into both oocytes and spermatogonia, respectively, depending on the particular gonadal environment they occur in (36–39). By contrast, we found no effect of the recipient’s gonadal environment on the ability or inability of transplanted germ cells to undergo clonal development. In particular, *C. elongatoides* male stem germ cells transplanted into ETT female recipients consistently developed into reduced oocytes that gave rise to recombining pure-bred progeny, while triploid male stem germ cells demonstrated the ability of PMER when transplanted into *C. elongatoides* females and produced purely clonal progeny, or tetraploid progeny with clonally transmitted maternal genome and incorporated sperm. This suggests that ability to undergo PMER is inherent to the hybrid constitution of the asexual gonial cells rather than being affected by their gonadal environment.

Our data also provided interesting insight into asymmetrical reproduction patterns of hybrids with respect to sex, since the hybrids’ spermatogonia transplanted into *C. elongatoides* males apparently didn’t express the ability of PMER and arrested their development in metaphase I due to abnormal pairing of orthologous chromosomes. These results are in exact agreement with previous analyses of natural and experimental ET and ETT hybrids (9, 40), which reported that PMER is confined to hybrid females and occurs already in the F1 generation, while all hybrid males are sterile with spermatogonia arrested at metaphase I (9, 30, 31, 41). How and why should the sex affect the initiation of PMER remains unclear, however. Our data demonstrated that gonial cells originating from male-determined juveniles are able to undergo PMER and develop into clonal progeny when transplanted into the ovary where they transdifferentiate into oogonia. This implies that in *Cobitis* hybrids, the ability of PMER reflects the phenotypic sex of gonads and does not necessarily depend on genetic sex determination. By contrast, investigation of hybrids from the related genus *Misgurnus*, (42, 43) demonstrated that female hybrids which were sex-reversed into males maintained the ability to produce unreduced fertile sperm *via* PMER, while natural hybrid males remained sterile (11). This would suggest that the cell’s capability of PMER may depend on genetic sex determination even when developing into male phenotypes.

The apparent discrepancy between our and Yoshikawa *et al*.’s (43) study may be due to several reasons. First, asexual hybrids in both genera evolved independently from different parental species and hence the particular type of cell deregulation leading to PMER may be different. Second, there are to date no robust data on genetic sex determination in Cobitidae, albeit male heterogamy (X1X2Y and X0, respectively) has been indicated in two species of the family (44, 45). Hence, the type of sex determination may vary between both *Cobitis* and *Misgurnus* genera. Finally, Yoshikawa *et al*. (43) investigated the development of female-originated gonial cells in phenotypic males, while we investigated the fate of male-originated gonial cells transplanted into recipients of both sexes. It is thus theoretically possible that hybrid primordial germ cells genetically determined as females maintain their capability of PMER even when turned into spermatogonia [as in (43)], while primordial germ cells genetically determined as males gain such a capability only when turned into female phenotypes, and produce sterile gametes when maintaining their original sex (as in present study).

Either way, our findings, together with previously gathered information about germ cell development in fishes, and in Cobitidae in particular, lead us to propose the following hypothesis related to the triggering of asexuality: the capability of clonal gametogenesis (at least the one based on PMER) is rather independent of the gonadal environment and appears triggered by intrinsic stimuli within asexual gonial cells, which is causally linked to the hybrid constitution of their genomes. Nonetheless, the very execution of PMER appears to be primarily bound to an oogonial developmental pathway. It is at this level when the gonadal environment affects development and asexuality, since primordial germ cells are sexually bipotent and their differentiation into oogonia is decisively affected by the environment in which they occur. Once the developmental pathway into male or female germlines is decided, the hybrid’s gonial cells develop into sterile spermatocytes, while the testes or fertile oocytes are capable of PMER while in the ovary. This hypothesis implies two sets of crucial questions for future research in asexual organisms: (1) How does the hybridization *per se* trigger PMER? and (2) Why is it usually linked to one sex in hybrids? With currently available knowledge, we may so far offer only speculative answers.

First, PMER occurs already in F1 generations (9, 25, 40) and hence it is unlikely that this trait evolves by accumulated mutations during evolution of hybrid populations. Instead, it is more likely that the execution of PMER is based on developmental programs that have already existed in cells of sexual progenitors of hybrid asexuals but are just triggered by the hybrid nature of the gamete. Possibly, the initiation of PMER is driven by accumulated incompatibilities between genomes brought together by hybridization, which fail to properly regulate gametic development and cell division leading to aberrant chromosome duplications (5, 6, 18). For instance, the very nature of PMER, *i*.*e*. multiplication of the genome without cell division, makes it at least superficially analogous to endopolyploidy, which is a common mechanism how various organisms, including fishes, modify the genomic content of specific cell types or tissues. Cellular mechanisms ensuring the alternation of S and G phases are relatively conserved among various animal lineages (46), suggesting that cells of most organisms are capable of endopolyploidy under the proper regulatory stimulus (47). Extrapolating Carman’s model (8) of asexuality, it is possible that PMER occurs in hybrid lineages when a particular type of misregulation between admixed parental subgenomes generates endopolyploidy specifically in gonial tissue, leading to stabilization of clonal lineage. The situation may be quite complex, however, and the result may crucially depend on other traits than hybridization, *e*.*g*. the hybrid’s ploidy affecting the stoichiometric ratio of orthologous alleles and their products. For instance, in *Poeciliopsis* spp. (mollies), diploid hybrids are hybridogenetic (*i*.*e*. clonally transmit only one parental genome and exclude the other’s before meiosis), while triploid hybrids between the same parental species change the reproductive mode to gynogenesis and clonality (48) Similarly, diploid hybrid *Misgurnus* spp. reproduce gynogenetically *via* PMER, while tetraploid hybrids between the same parental species produce reduced gametes (30)

The second question may have a lot in common to fundamental differences between male and female types of gametogenesis. Such differences may translate into the timing of DNA methylation in male and female gametogenesis (49, 50). There is also evidence for differences in patterns of epigenetic regulation between SSCs and derived oocytes from SSCs (51), which suggest an artificial epigenetic restart of our transplanted SSCs. It may thus be proposed that in the hybrid’s spermatogonia transplanted into female recipients, the cell-cell communication between female somatic cells and transplanted SSCs has led to the establishment of gonadal tissue according to the recipient’s sex determination (52), thus epigenetic reprogramming to female-like patterns and ultimate awakening of PMER. In that scenario, the SSCs transplanted into male recipients would not undergo such a process. They would thus not gain the ability of PMER.

The present findings demonstrated that the investigation of gametic development is likely to provide crucial insights in understanding asexual reproduction and the establishment of interspecific reproductive barriers in the speciation process. Namely, this study indicated that ability to perform asexual gametogenesis *via* PMER is causally linked to hybrid composition gonial cells and is triggered by factors intrinsic to these cells and developmental programs inherited from parental species. On the other hand, it also appears that the execution of PMER is exclusive to the female germline, whose determination apparently depends on cell-cell communication with surrounding gonadal tissue. Thus, even in hybrid females, whose fertility is restored by PMER, the sex-specific factors of surrounding somatic tissue that control gametic development contribute to the postzygotic barrier, since PMER prevents the hybrid’s effective backcrossing to parental species.

## Materials and Methods

Experimental protocol was approved by Ministry of Agriculture of the Czech Republic (reference number: 55187/2016-MZE-17214).

### Sterilization by *dead end* antisense oligonucleotide

Morpholino oligonucleotide was designed based on the *dnd* gene sequence of *Misgurnus anguillicaudatus* (AB531494.1). The *dnd* sequence of *Cobitis taenia* obtained from DDBJ/EMBL/GenBank transcriptome (GGJF00000000.1) was aligned against AB531494.1 by muscle software v.3.8.1551 to validate specificity to the *Cobitis* genus. Alignment with the highlighted MO target and phylogenetic maximum likelihood tree was performed with iqtree, 1.6.10 (53) and approximate Bayes test (54). MFP+MERGE model selection was based on the *dnd* gene of *M. anguillicaudatus* (AB531494.1), *Danio rerio* (AY225448.1), *Carrasius auratus* (JN578697.1:44-1140), *Gobiocypris rarus* (KM044011.1), *Paedocypris progenetica* (KY828447.1), *Sinocyclocheilus rhinocerous* (XM_016576295.1), *Rhodeus ocellatus* (MG995743.1). SNPs were checked at probe position for any interspecific variability. Morpholino oligonucleotide was synthesized by Gene Tools, LLCTM (Philomath, Oregon, U.S.A.).

The final solution for sterility induction was composed of 100 μM of MO and 300 ng/μL of mRNA in combination with GFP and zebrafish (*D. rerio*) nos1 3’UTR and diluted in 0.2 M KCl (55). The control group solution received only 300 ng/μL of mRNA diluted in 0.2 KCl. Solutions were loaded into a microcapillary mounted on a micromanipulator (M-152 Narishige, Japan) with an automatic microinjector (FemtoJet Eppendorf, Germany). Each embryo was injected into a blastodisc at the one to four cell stage. Altogether, 50 embryos were injected in each group.

### Histology analysis

Either whole body segments or gonadal tissue were fixed overnight in Bouin’s fixative. Specimens were immersed in 70% ethanol, dehydrated and cleared in an ethanol–xylene series, embedded into paraffin blocks, and cut transversally into 4 μm thick sections using a rotary microtome (Leica RM2235; Wetzlar, Germany). Paraffin slides were stained with hematoxylin and eosin by using a staining machine (Tissue-Tek DRS 2000; Sakura Finetek USA, Inc., Torrance, California) according to standard procedures. Histological sections were photographed using a microscope (Nikon Eclipse Ci; Tokyo, Japan) with a mounted camera (Canon EOS 1000D; Ōta, Tokyo, Japan). In the case of morpholino treated fish, the sex identification was based on Fujimoto *et al*. (34) and Goto *et al*. (56).

### Induction of germline chimeras and donor-derived gametes production

Five diploids *C. elongatoides* and two triploids *C. elongatoides-taenia-taenie* donor male specimens were over anesthetized in tricaine solution (MS222), disinfected with 70% ethanol and decapitated. The body cavity was carefully opened, and the gonads were removed and placed in ice-cold phosphate-buffered saline (PBS). Testes were cut into smaller fragments to allow leakage of the sperm and serially washed in PBS. Gonad fragments were transferred into 15 ml tubes and well chopped with scissors. Gonadal tissue was enzymatically digested in 5 ml of PBS with 0.15% trypsin with a laboratory shaker at room temperature for 1.5 h. DNase I (Sigma Aldrich 10104159001; Merck, Burlington, Massachusetts, USA) (aliquoted to 5% stock solution in RNase free water) was added continuously when clumping was observed. Afterwards, digestion was terminated by the addition of 5 ml L15 with 20% fetal bovine serum, filtrated through 30μm filters, and centrifuged at 400*g* for 10 min. The supernatant was removed, and the pellet was carefully resuspended. The cell suspension was loaded into a pulled glass microcapillary mounted on a micromanipulator with a pneumatic injector.

Prior to cell transplantation, primordial germ cells were depleted and, 2n and 3n recipients were prepared by injection of antisense morpholino designed against the *dead-end* gene. Sterilized recipients were anesthetized in 0.05% tricaine solution, placed on an agar coated Petri dish and cells were injected into the coelomic cavity. Recipients were transferred into fresh water to recover. Germline chimeras were cultured at room temperature in aquaria with controlled cooling and water filtration and fed *ad libitum* with brine shrimps (*Artemia* sp.), bloodworms (*Tubifex* sp.), and a dry diet. Two groups of chimeras were made: 1) diploid recipient and triploid male donor (DrTd) and 2) triploid recipient and diploid male donor (TrDd).

### Spawning

Six months prior to spawning, water temperature was slowly decreased (−2 °C per day) to 14 °C and kept at this level for 3 months. In the following 3 months, water temperature was increased (+2 °C per day) to final temperature of 22 °C and kept till spawning with increased bloodworm feeding. Transplanted fish were exactly 2 years old at the time of spawning. Transplanted fish together with control fish (*C. elongatoides*) were injected twice (24 and 12 h prior to spawning) with Ovopel (Interfish Kf, Budapest, Hungary). A solution for the first injection was made from one Ovopel pill per 20 mL of 0.9% NaCl. The solution for the second injection was made from one Ovopel pill per 5 mL of 0.9% NaCl. In both cases, the volume of the Ovopel solution directly injected into a fish body was 0.05 mL per 10 g of fish weight. Eggs were carefully removed from spined loaches and put in a dried Petri dish. Sperm were added to the eggs together with fresh water. The number of fish in each group was as follows: 14 fish in the DrTd group (six females, eight males), eight fish in TrDd (eight females, zero males), and six males of *C. elongatoides* as controls.

### Identification of chimeras

In this study, we used parental species *C. elongatoides* (EE – composition of genomes) and triploid hybrids between *C. elongatoides and C. taenia* (ETT). Diploid parental species produce haploid gametes while hybrids produce oocytes which contain both genomes; therefore presence/absence of C. *taenia* (TT) species in offspring can prove the success of transplantation. Successfully transplanted 2n EE fish will produce ETT eggs while 3n ETT fish will produce haploid eggs with genome E. Indication of five specific loci Cota068, Cota111, Cota010, Cota093, Cota032 of *C. taenia* (57) in offspring from parental species of EE means that diploid fish possess hybrid gonads. On the other hand, triploid fish ETT producing haploid eggs represent the occurrence of diploid gonads in triploid fish. To support our results, we also used flow cytometry analysis and meiosis analysis on both parental species and offspring.

### DNA extraction and analysis of microsatellites

Whole genomic DNA from individuals tested (*C. elongatoides* and *C. elongatoides-taenia-taenia*) were extracted from a dorsal fin in adults or part of larvae using a commercial Tissue DNA Isolation Kit (Geneaid Biotech, Taipei, Taiwan) following the manufacturer’s protocol. Genotype determination in the fishes was performed by analyses of selected microsatellite species specific loci (31, 57). Fragment-length analyses were performed on an ABI 3730 Avant capillary sequencer (Applied Biosystems, Foster City, California, USA) with an internal size standard (GeneScan-500 LIZ, Thermo Fisher Scientific, Waltham, Massachusetts, USA); the alleles were scored manually with GeneMapper v. 3. 7 (Applied Biosystems, Zug, Switzerland).

### Flow cytometry analysis

The level of ploidy was determined as the relative DNA content of fin clip cells *via* flow cytometry (Partec CCA I; Partec GmbH, Munster, Germany with a UV mercury lamp for excitation and an emission level of 435/500 nm) using standard CyStain^®^ DNA 1-step solution (Sysmex CZ s.r.o., Brno, Czech Republic) containing 49.6-diamidino-2-phenylindol (DAPI). As a reference standard, we used a fin clip of diploid *C. elongatoides*.

### Pachytene chromosomes with immunofluorescent staining

Pachytene chromosomes were obtained from males and females according to protocols described by Moens (58) and Araya-Jaime *et al*. (59). Ovaries were homogenized manually in 1× PBS solution. Afterwards, 20 μl of cells suspension was put on SuperFrost^®^ slides (Menzel Gläser; Thermo Fisher Scientific) followed by addition of 40 μl of 0.2 M sucrose and 40 μl of 0.2% Triron X100 for 7 min. The samples were fixed for 16 min by adding 400 μl of 2% PFA. Testes were homogenized manually followed by dropping 1 μl of suspension into 30 μl of hypotonic solution (1/3 of 1× PBS) and then dropped onto SuperFrost^®^ slides (Menzel Gläser; Thermo Fisher Scientific). The samples were fixed in 400 μl of 2% PFA for 4 min. After fixation, slides with the pachytene samples from males and females were air dried and washed in 1× PBS.

Slides were stored until immunofluorescent staining of synaptonemal complexes (SC). Lateral components of synaptonemal complexes (SCs) were visualized by rabbit polyclonal antibodies (ab14206, Abcam) against SYCP3 protein while the central component of SCs was detected by chicken polyclonal antibodies against SYCP1 protein (a gift from Sean M. Burgess). Fresh slides were incubating with 1% blocking reagent (Roche) in 1× PBS and 0.01% Tween-20 for 20 min followed by the addition of primary antibody for 1h at RT. Slides were washed 3 times in 1× PBS at RT and incubated in the combination with secondary antibodies Alexa 488-conjugated goat anti-rabbit IgG (H+L) (Molecular Probes) and Alexa-594-conjugated goat anti-chicken IgG (H+L) (Molecular Probes) for 1h at RT. Slides were washed in 1× PBS and mounted in Vectashield/DAPI (1.5 mg/ml) (Vector, Burlingame, Calif., USA).

Diplotene chromosomal samples (also known as “lampbrush chromosomes”) were prepared from parental and hybrid females according to an earlier published protocol (60). Vitellogenetic oocytes of 0.5– 1.5 mm in diameter were taken from females in the OR2 saline [82.5 mM NaCl, 2.5 mM KCl, 1 mM MgCl2, 1 mM CaCl2,1mM Na2HPO4, 5 mM HEPES (4-(2-hydroxyethyl)-1-piperazineethanesulfonic acid); pH 7.4]. Isolation of the oocytes’ nuclei was performed manually in the isolation medium “5:1” (83 mM KCl, 17 mM NaCl, 6.5 mM Na2HPO4, 3.5 mM KH2PO4, 1mM MgCl2, 1 mM DTT (dithiothreitol); pH 7.0–7.2) using jeweler forceps (Dumont, Switzerland). Nuclear envelopes were manually removed in a quarter strength “5:1” medium with the addition of 0.1% paraformaldehyde and 0.01% 1M MgCl2 in glass chambers attached to a slide. After this procedure, we obtained chromosome samples from individual oocytes in each chamber. Slides with oocyte nuclei contents were subsequently centrifuged for 20 min at +4 °C, 4000 rpm, fixed for 30 min in 2% paraformaldehyde in 1x PBS, and post-fixed in 70% ethanol overnight (at +4°C).

Pachytene and diplotene chromosomes were investigated using a Provis AX70 Olympus microscope with standard fluorescence filter sets. Microphotographs were captured by CCD camera (DP30W Olympus; Tokyo, Japan). Olympus Acquisition Software was used for capturing the images followed by their adjustment and arrangement in Adobe Photoshop, CS6 software.

## Acknowledgments

This research was funded by Institutional Research Concept RVO67985904 and by Czech Science Foundation, projects No. 17-09807S (to M.P. & K.H.) and 19-21552S & 21-25185S (to Lab. of Fish Genetics). The transplantation and fish manipulation were financially supported by the Ministry of Education, Youth and Sports of the Czech Republic - project CENAKVA (LM2018099) and Biodiversity (CZ.02.1.01/0.0/0.0/16_025/0007370).

Prof. Sean Burgess for sharing primary antibodies against SYCP1.

